# Prolyl Endopeptidase (PREP) is Involved in the Reproductive Functions and Cytoskeletal Organization in Rat Spermatogenesis and in Mammalian Sperm

**DOI:** 10.1101/319491

**Authors:** Massimo Venditti, Sergio Minucci

**Affiliations:** Dipartimento di Medicina Sperimentale – Sez. Fisiologia Umana e Funzioni Biologiche Integrate “F. Bottazzi” – Università degli Studi della Campania “Luigi Vanvitelli”, via Costantinopoli 16 – 80138 Napoli – Italy

**Keywords:** PREP, Tubulin, Testis, First wave of spermatogenesis, Spermatozoa

## Abstract

Prolyl endopeptidase (PREP) is an enzyme which cleaves several peptide hormones and neuropeptides at the carboxyl side of proline residues, involved in many biological processes, including cell proliferation and differentiation, glucose metabolism, learning, memory and cognitive disorders. Moreover, PREP was identified as binding partner of tubulin, suggesting that this endopeptidase may be involved in microtubule-associate processes, independent of its peptidase activity. Several reports have also suggested PREP participation in both male and female reproduction-associated processes. In this work, we assessed the possible association of PREP with the morphogenesis of rat testis, profiling its localization versus tubulin, during the first wave of spermatogenesis and in the adult gonad (from 7 to 60 dpp). Here we show that, in mitotic phases, PREP shares its localization with tubulin in Sertoli cells, gonocytes and spermatogonia. Later, during meiosis, both proteins are found in spermatocytes, and in the cytoplasm of Sertoli cells protrusions, which surround the germ cells, while, during spermiogenesis, they both localize in the cytoplasm of round and elongating spermatids. Finally, they are expressed in the flagellum of mature gametes, as corroborated by additional immunolocalization analysis on both rat and human sperm. Our data strongly support the hypothesis of a role of PREP in supporting a correct reproductive function and in cytoskeletal organization during Mammalian testis morphogenesis and gamete progression, while also hinting at its possible investigation as a morphological marker of germ cell and sperm physiology.

**Summary statement:** In this paper we show the co-localization of the enzyme PREP with tubulin during the first wave of rat spermatogenesis and in mature gametes of rat and human.

## Introduction

Prolyl endopeptidase (PREP; EC 3.4.21.26) is a protein belonging to the serine protease family, widely conserved through evolution (Venäläinen et al., 2004). It was identified for the first time in the human uterus (Walter et al. 1971), but soon detected in all mammalian tissues, including liver, kidney, heart, spleen, and brain, where it shows the highest enzymatic activity (Yoshimoto et al., 1979; Taylor et al., 1980). PREP has a typical endopeptidase structure, including the catalytic triad formed by Ser554, Asp641 and His680 (Rea and Fülöp, 2006). PREP is able to hydrolyze the peptide bond on the carboxyl side of proline residues in oligopeptides comprising no more than about 30 amino acid residues (Szeltner and Polgár, 2008), as well as peptide hormones and neuropeptides (Mentlein, 1988; Wilk, 1983). Despite its common cytosolic localization and the lacks of a secretion signal or a lipid anchor sequence (Venäläinen et al, 2004), it is believed that PREP may be released from the cells and act outside by inactivating extracellular neuropeptides (Ahmed et al., 2005). PREP has been implicated in several biological processes, including cell proliferation and differentiation (Matsubara et al., 1998; Suzuki et al., 2014), cell death (Bär et al., 2006; Matsuda et al., 2013), glucose metabolism (Kim et al., 2014), celiac disease (Siegel et al., 2006; Comino et al., 2013), learning and memory (Irazusta et al., 2002; D’Agostino et al., 2013) and cognitive disorders (Rossner et al., 2005; Hannula et al., 2013). Further reports about the intracellular activity of PREP suggested an additional physiological role for this enzyme (Schulz et al., 2005). Indeed, PREP was identified as binding partner of tubulin, indicating novel functions for PREP in vesicle transport and protein secretion (Morawski et al., 2011). As well known, microtubules are highly dynamic cytoskeletal components that play fundamental roles in many cellular processes, such as motility, intracellular transport, division and cell shape (Jordan and Wilson, 2004; Conde and Cáceres, 2009; Helmke et al., 2013). Since cytoskeletal remodeling is a critical feature which allows the cell to modulate its shape and architecture, the study of its actors during gametogenesis and reproduction is of great interest, as the germinal compartment and the germ cells undergo a complex series of transformations throughout the process (Venditti and Minucci, 2017), led by heavy cytoskeletal elements organization (Lie et al., 2010). So far, only a few reports have already suggested PREP participation in both male and female reproduction-associated processes (Kimura et al., 1998; Kimura et al., 2002; Dotolo et al., 2016). Thus, in this work, we assessed the possible association of PREP with the morphogenesis of rat testis, by studying and comparing its expression and localization with tubulin, during the first wave of spermatogenesis and in the adult tissue. We also extended our profile to rat and human spermatozoa, in order to further enhance such profile and to clarify PREP distribution in mature gametes.

## Materials and Methods

### Animal care, tissue extraction, and collection of rat spermatozoa

Male Sprague–Dawley rats (*Rattus norvegicus*) were housed under definite conditions (12D:12L) and they were fed with standard food and provided with water *ad libitum*. Animals at different development stages (7 days post-partum, 14, 21, 28, 35, 42, 60 dpp, and adult) were sacrificed by decapitation under Ketamine anaesthesia (100 mg/kg i.p.) in accordance with national and local guidelines covering experimental animals. For each animal testes were dissected; one testis was fixed in Bouin’s fluid and embedded in paraffin for histological analysis, one was quickly frozen by immersion in liquid nitrogen and stored at −80°C until protein extraction. Additionally, epididymides were removed from adult rats and minced in phosphate buffer saline, PBS (13.6 mM NaCl; 2.68 mM KCl; 8.08 mM Na2HPO4; 18.4 mM KH2PO4; 0.9 mM CaCl2; 0.5 mM MgCl2; pH 7.4) to let the spermatozoa (SPZ) flow out from the ducts. Then, the fluid samples were filtered and examined under a light microscope to exclude contamination by other cell types. Next, aliquots were spotted and air-dried on slides, then stored at - 20°C, while the remaining samples were centrifuged at 1,000g for 15 min at 4°C and stored at - 80°C until protein extraction.

### Collection of human spermatozoa

Human sperm from qualified donors was centrifuged at 800g for 10 min; the supernatant was removed and the pellet was washed and resuspended in PBS. The samples were examined under a light microscope and aliquots were spotted and air-dried on slides, then stored at - 20°C, while the remaining samples were centrifuged at 1,000g for 15 min at 4°C and stored at 80°C until protein extraction.

### Preparation of total protein extracts and Western blot analysis

The testes and SPZ (from rat or human) were lysed in a specific buffer (1% NP-40, 0.1% SDS, 100 mM sodium ortovanadate, 0.5% sodium deoxycholate in PBS) in the presence of protease inhibitors (4 mg/ml of leupeptin, aprotinin, pepstatin A, chymostatin, PMSF, and 5 mg/ml of TPCK). The homogenates were sonicated twice by three strokes (20 Hz for 20 s each); after centrifugation for 30 min at 10,000g, the supernatants were stored at - 80°C. Proteins from testis and SPZ (50 µg) were separated by 9% SDS-PAGE and transferred to Hybond-P polyvinylidene difluoride (PVDF) membranes (Amersham Pharmacia Biotech, Buckinghamshire, UK) at 280 mA for 2.5 h at 4°C. The filters were treated for 3 h with blocking solution [5% skim milk in TBS (10 mM Tris–HCl pH 7.6, 150mM NaCl)] containing 0.25% Tween-20 (Sigma–Aldrich Corp., Milan, Italy) before the addition of anti-PREP (Abcam Cat #ab58988), or anti-Tubulin (Sigma–Aldrich Corp., Milan, Italy) antibody diluted 1: 5,000 and 1: 10,000 respectively, and incubated overnight at 4°C. After three washes in TBST (TBS including 0.1% Tween20), the filters were incubated with horseradish peroxidase-conjugated anti-rabbit IgG (Sigma–Aldrich Corp., Milan, Italy) for the rabbit anti-PREP antibody, or anti-mouse IgG (Sigma–Aldrich Corp., Milan, Italy) for the mouse anti-Tubulin antibody, both diluted 1: 10,000 in the blocking solution. Then, the filters were washed again three times in TBST and the immunocomplexes were revealed using the ECL-Western blotting detection system (Amersham Pharmacia Biotech).

### Tissue quality control and classification of testicular cell types

In order to assess the quality of the tissue samples and their staging, 7 mm-thick rat testis sections of all samples (7, 14, 21, 28, 35, 42, 60 dpp) were prepared and a haematoxylin-eosin staining was performed (see Fig. 1). The cell types for each time point were characterized and confirmed following previously reported classifications (Picut et al., 2015; Pariante et al., 2016).

**Fig. 1.**
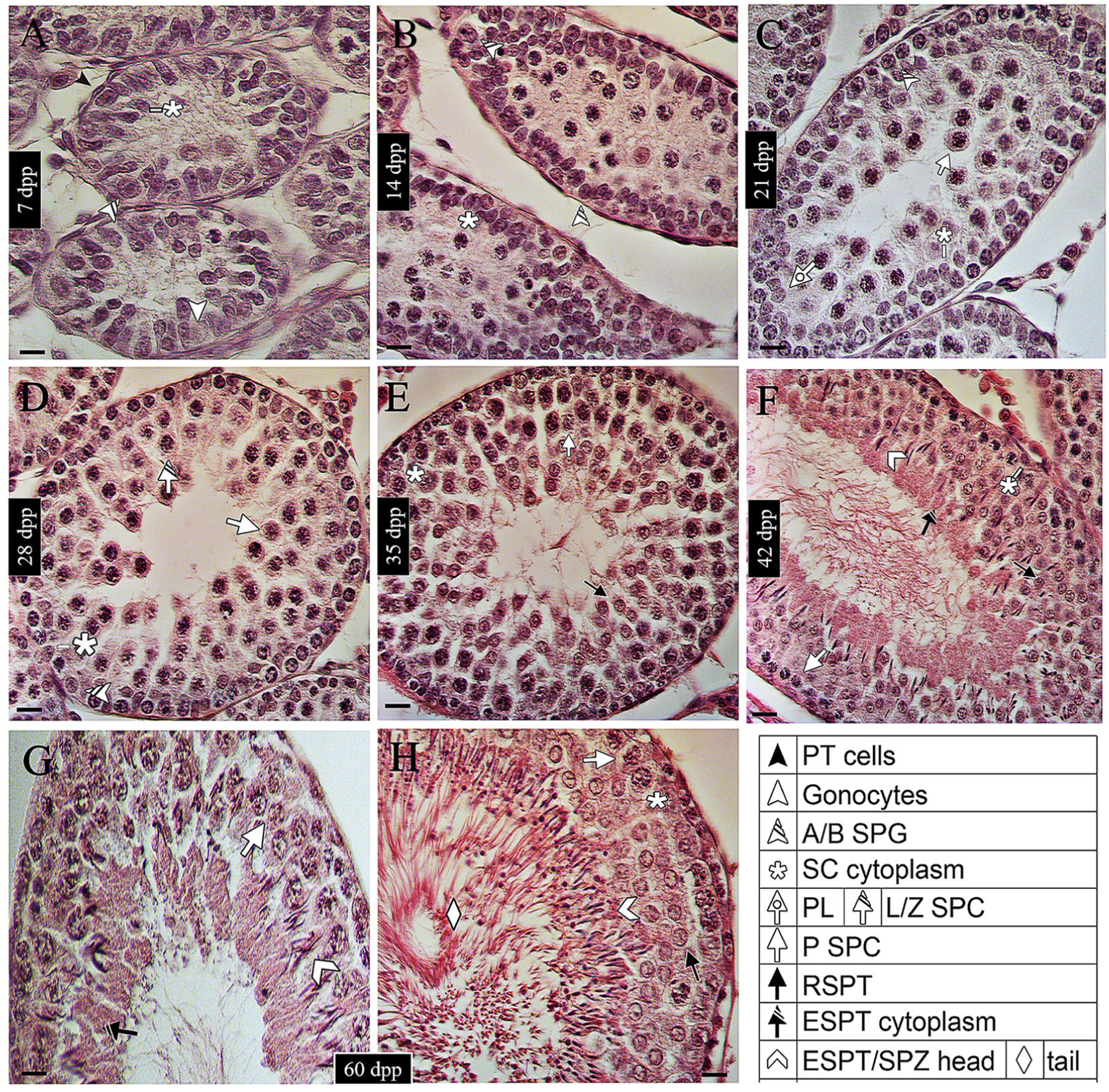
Histology and staging of the developing rat testis. Haematoxylin-eosin staining of tissue sections at 7 (A), 14 (B), 21 (C), 28 (D), 35 (E), 42 (F), 60 (G and H) dpp, in which the most representative cell types are highlighted (for review see Picut et al., 2014). Pointer legend is provided in the bottom-right table. PT cells: Peritubular cells; SPG: Spermatogonia; SC: Sertoli cells; PL SPC: Pre-leptotene primary Spermatocytes; L/Z: Leptotene/Zygotene; P: Pachytene; RSPT: Round Spermatids; ESPT: Elongating Spermatids; SPZ: Spermatozoa. Scale bars represent 20 μm.

### Immunofluorescence analysis on rat testis

For PREP co-localization with both Tubulin and the acrosome system, 7 mm-testis sections were dewaxed, rehydrated, and processed as described by Venditti et al. (2018). Antigen retrieval was performed by pressure cooking slides for 3 min in 0.01 M citrate buffer (pH 6.0). Then, the slides were incubated with 0.1% (v/v) Triton X-100 in PBS for 30 min. Later, nonspecific binding sites were blocked with an appropriate normal serum diluted 1:5 in PBS containing 5% (w/v) BSA before the addition of anti-PREP, or anti-Tubulin antibody diluted 1:100, for overnight incubation at 4°C. After washing in PBS, slides were incubated for 1 h with the appropriate secondary antibody (Anti-Rabbit Alexa Fluor 488, Invitrogen; FITC-Jackson, ImmunoResearch, Pero MI, Italy; Anti-Mouse IgG 568, Sigma-Aldrich, Milan, Italy) diluted 1:500 in the blocking mixture and with PNA lectin (Alexa Fluor 568, Invitrogen, Monza MB, Italy) diluted 1:50. The slides were mounted with Vectashield + DAPI (Vector Laboratories, Peterborought, UK) for nuclear staining, and then observed with a microscope then observed under the optical microscope (Leica DM 5000 B + CTR 5000) and images where viewed and saved with IM 1000.

### Immunofluorescence analysis on SPZ

To determine PREP and Tubulin co-localization in rat and human SPZ, the samples were firstly fixed in 4% paraformaldehyde in PBS, and then washed in phosphate buffer (0.01 M PBS, pH 7.4). The slides were incubated with 0.1% (v/v) Triton X-100 in PBS for 30 min. Later, nonspecific binding sites were blocked with an appropriate normal serum diluted 1:5 in PBS containing 5% (w/v) BSA before the addition of the primary antibody (PREP and Tubulin), as described above, and overnight incubation at 4°C. After washing in PBS, slides were incubated for 1 h with the appropriate secondary antibody (Anti-Rabbit Alexa Fluor 488, Invitrogen; FITC-Jackson, ImmunoResearch, Pero MI, Italy; Anti-Mouse IgG 568, Sigma-Aldrich, Milan, Italy) diluted 1:500 in the blocking mixture and with PNA lectin (Alexa Fluor 568, Invitrogen, Monza MB, Italy) diluted 1:50. The slides were mounted with Vectashield + DAPI (Vector Laboratories) for nuclear staining, then observed under the optical microscope (Leica DM 5000 B + CTR 5000) with UV lamp, and images where viewed and saved with IM 1000.

## Results

### Expression of PREP and Tubulin during the post-natal development of rat testis

The expression of PREP during the postnatal development of the gonad was assessed by Western Blot analysis on protein extracts from some of the most representative time points during the first wave of spermatogenesis: 7 dpp (transition of gonocytes from tubule lumen toward the base; 14 dpp (proliferation of Sertoli cells and A and B spermatogonia, before meiosis); 21 dpp (presence of spermatocytes, which undertake meiosis; first phases of blood-testis barrier formation); 28 dpp (conclusion of “first wave” meiosis, completion of the blood-testis barrier); 35 dpp (presence of newly-formed round spermatids in spermiohistogenesis); 42 dpp (final steps of spermiohistogenesis); 60 dpp (mature testis; presence of spermatozoa and of all the characteristic germ cell associations). A band of the expected size (80 kDa) was detected for PREP in all samples (Fig. 2). The same time-point progression was employed for the analysis of Tubulin. As expected, bands were detected in all samples, as a confirmation of the expression of this cytoskeletal protein during testis development (Fig. 2).

**Fig. 2.**
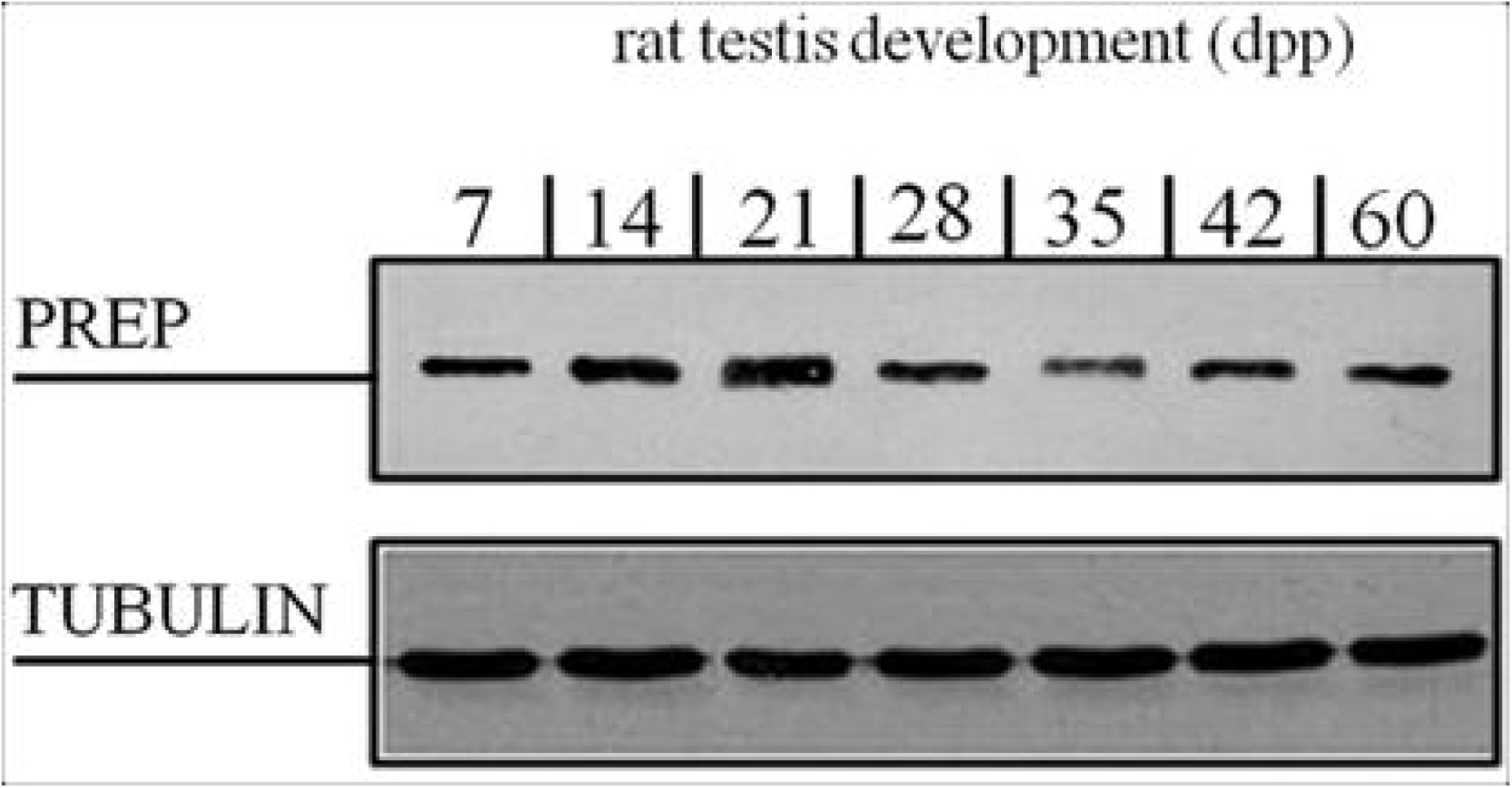
Expression of PREP and Tubulin during the post-natal development of rat testis. Western blot analysis which shows the expression of PREP (81 KDa, top section) and Tubulin (50 KDa, bottom section) during rat post-natal development, at 7,14,21,28,35,42, and 60 days post-partum (dpp). The two proteins are always expressed.

### Localization of PREP during the post-natal development of rat testis

First, tissue quality and staging were checked by performing a haematoxylin-eosin staining on sections of rat testis at the same time points as described in the previous paragraph (Fig. 1). PREP localization was studied by immunofluorescence analysis on developing testis sections (7, 14, 21, 28 dpp, Fig. 3; 35, 42, and 60 dpp, Fig. 4). At 7 dpp (Fig. 3 A-C), the protein signal was localized in Sertoli cell (SC) cytoplasm, but also detectable in luminal gonocytes (Fig. 3 B, C) and peritubular cells; at 14 dpp (Fig. 3 D-F), the signal was still localized in SC, and it was evident in A and B spermatogonia (SPG; Fig. 3 E-F). At 21–28 dpp (Fig. 3 G-L) it was detectable inside the cytoplasm of meiotic I spermatocytes (SPC; Fig. 3 H, I, K, L, better highlighted by the insets), as well as SC. During spermiogenesis, as shown from 28 dpp onward, it was possible to highlight the occurring acrosome formation, thanks to PNA lectin staining (Fig. 3 J-L, P; Fig. 3). In 35 dpp tubules (Fig. 4 A-C) PREP signal was detectable in the cytoplasm of SC, which extends from the base to the lumen, surrounding the germ cells (Fig. 4 A-C and insets). Then, the protein localized in elongating SPT at 42 dpp and was also detectable SC cytoplasm (Fig. 4 D-F). Finally, at 60 dpp (Fig. 4 G-I), after the conclusion of the first spermatogenetic wave, the signal was comparable to the one seen at 42 dpp, with the protein present in elongating spermatids and SC cytoplasm.

**Fig. 3.**
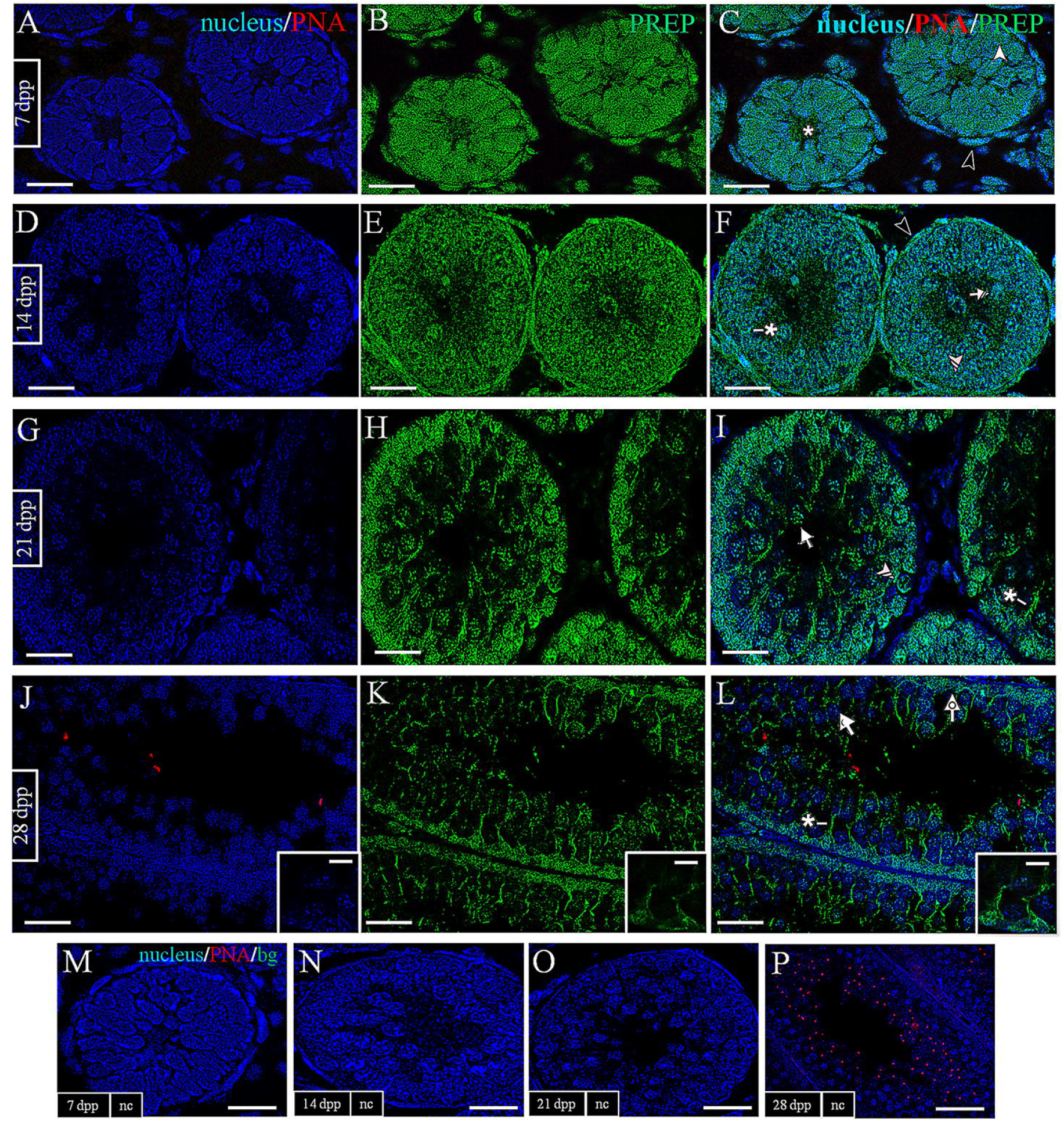
Localization of PREP during the post-natal development of rat testis, part 1 (7–28 dpp). A, D, G, J. DAPI-fluorescent nuclear staining (blue) and PNA lectin acrosome staining (red). B, E, H, K. PREP fluorescence (green). C, F, I, L, M, N, O, P. Merged fluorescent channels (blue/red/green). A, B, C. 7 dpp testis; PREP-positive fluorescence is detectable in the central region of the maturing tubules, as well as in the Sertoli cells cytoplasm. D, E, F. 14 dpp; fluorescent signal is present in spermatogonia and Sertoli cells. G, H, I. 21 dpp; J, K, L. 28 dpp; positive cells now include meiotic spermatocytes; Sertoli cells are not positive; insets show primary spermatocytes at different stages. At 28 dpp the acrosome formation starts to be visible through PNA-lectin staining. M, N, O, P. Negative controls for the same time points, obtained by omitting the primary antibody. Scale bars represent 20 μm, except for the insets, where they represent 10 μm. PNA: PNA lectin staining. bg: Background/autofluorescence. nc: Negative controls. For cell-type pointer legend, see table in Fig. 1.

**Fig. 4.**
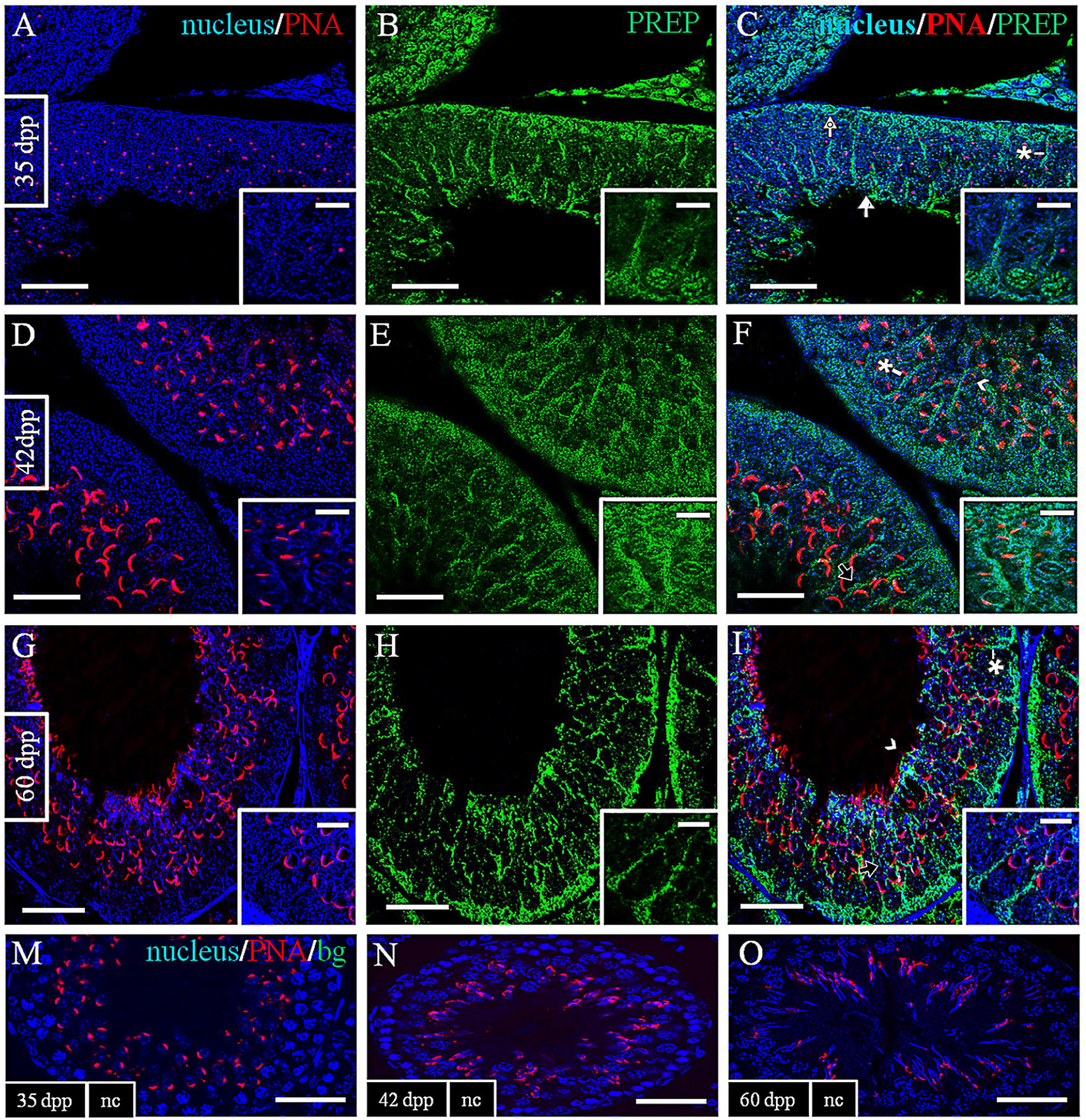
Localization of PREP during the post-natal development of rat testis, part 2 (35–60 dpp). A, D, G. DAPI-fluorescent nuclear staining (blue) and PNA lectin acrosome staining (red). B, E, H. PREP fluorescence (green). C, F, I, J, K, L. Merged fluorescent channels (blue/red/green). A, B, C. 35 dpp testis; D, E, F. 42 dpp; DAAM1 is detectable in spermatocytes and spermatids, where the acrosome signal is evident. G, H, I. 60 dpp; all cell types are positive; the cytoplasmic droplet is especially notable; insets show round spermatids (35 dpp), elongating spermatids (42 dpp) and spermatozoa (60 dpp). J, K, L. Negative controls for the same time points, obtained by omitting the primary antibody. Scale bars represent 20 μm, except for the insets, where they represent 10 μm. PNA: PNA lectin staining. bg: Background/autofluorescence. nc: Negative controls. For cell-type pointer legend, see table in Fig. 1.

### Co-localization of PREP and Tubulin during the post-natal development of rat testis

Given PREP association with Tubulin, the co-localization profile of the two proteins was performed on the same time-point described above. The immunofluorescence analysis showed that Tubulin signals resulted in a pattern comparable with PREP localization: Tubulin (Fig.5 and 6) was expressed in all stages and, to varying extent, by all cell types, but it was especially represented inside the somatic SC which nurse the mitotic and meiotic cells during the first phases of spermatogenesis (Fig. 5), as well as the SPT during their differentiation into SPZ (Fig. 6). PREP and Tubulin initially co-localize within GC junctions (Fig. 5 C, F and I). Form 28 dpp on they both are present inside SC cytoplasm, which surrounds the developing GC (Fig. 5 L, and Fig. 8 C, F), as well as, during spermiohistogenesis, in SPT, and in the epithelial cells which rearrange their architecture to support the path of the evolving germ cells (GC) toward the lumen (Fig. 6 F and I).

**Fig. 5.**
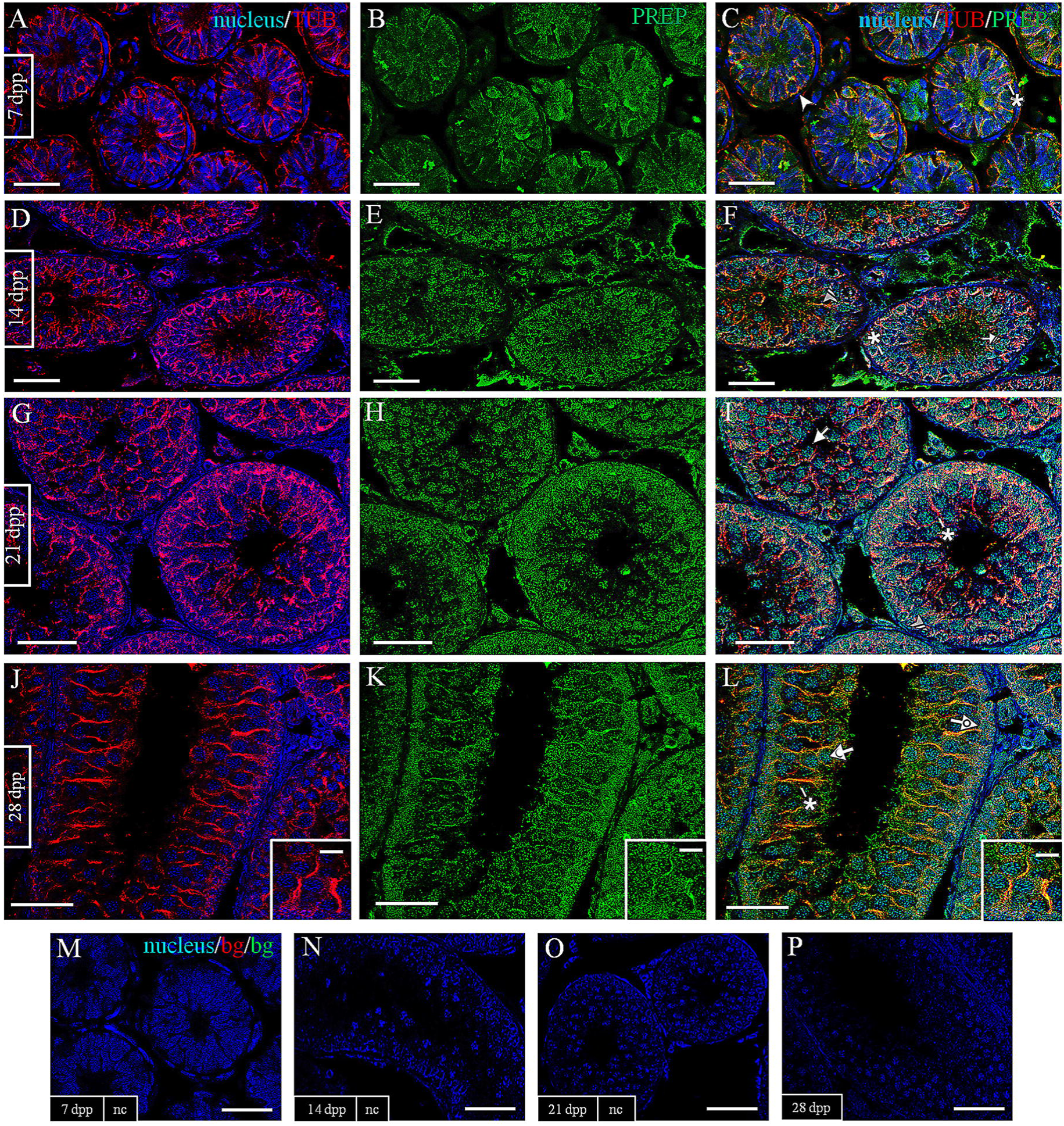
Co-localization of PREP and Tubulin during the post-natal development of rat testis, part 1 (7–28 dpp). A, D, G, J. DAPI-fluorescent nuclear staining (blue) and Tubulin staining (red). B, E, H, K. PREP fluorescence (green). C, F, I, L, M, N, O, P. Merged fluorescent channels (blue/red/green). A, B, C. 7 dpp testis; D, E, F. 14 dpp; G, H, I. 21 dpp; J, K, L. 28 dpp. Tubulin signal is strong in all samples, especially in the cytoplasm of Sertoli cells. M, N, O, P. Negative controls for the same time points, obtained by omitting the primary antibody. Scale bars represent 20 μm. TUB: Tubulin. bg: Background/autofluorescence. nc: Negative controls. For cell-type pointer legend, see table in Fig. 1.

**Fig. 6.**
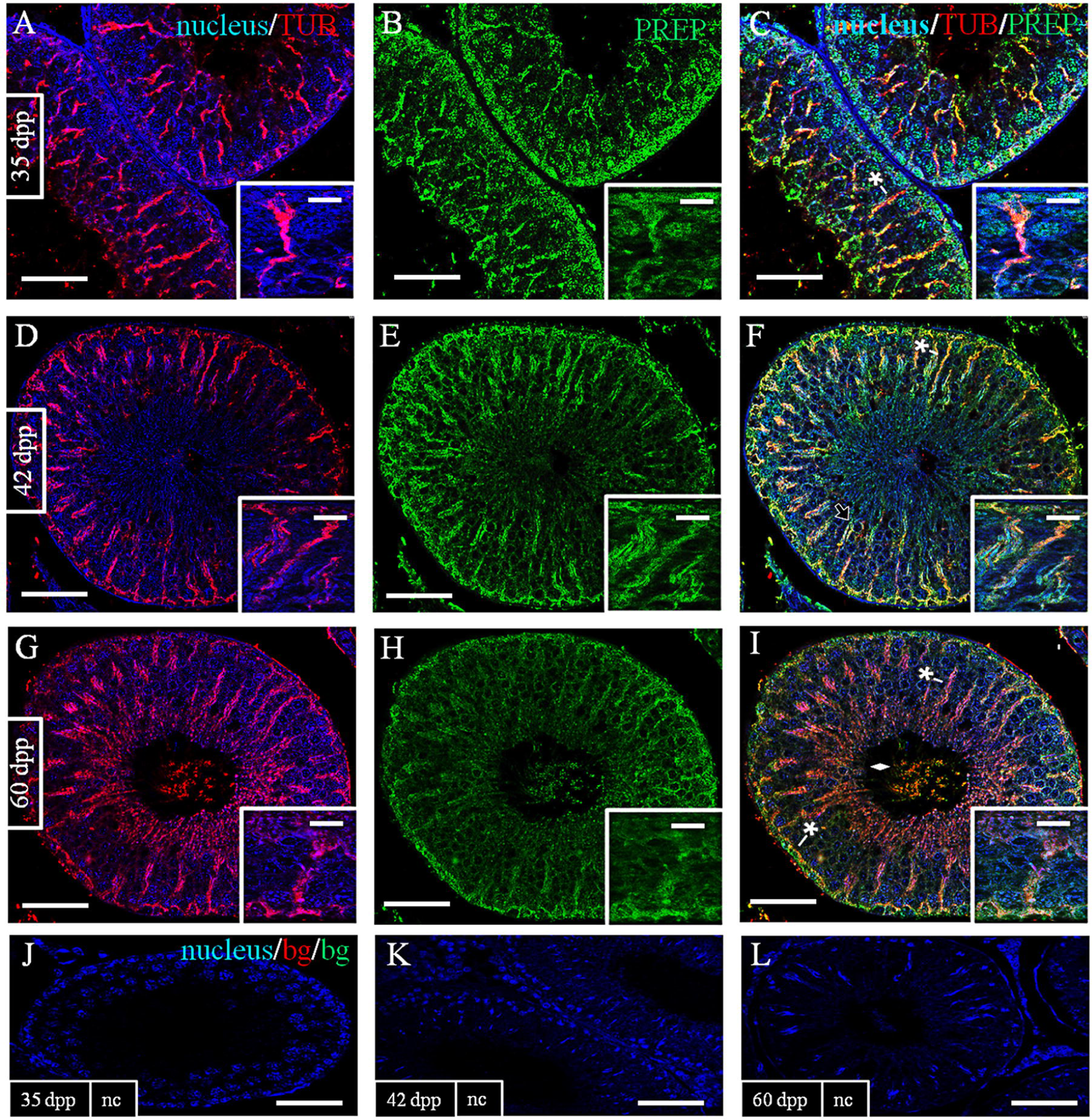
Co-localization of PREP and Tubulin during the post-natal development of rat testis, part 2 (35–60 dpp). A, D, G. DAPI-fluorescent nuclear staining (blue) and Tubulin staining (red). B, E, H. PREP fluorescence (green). C, F, I, J, K, L. Merged fluorescent channels (blue/red/green). A, B, C. 35 dpp testis; D, E, F. 42 dpp; G, H, I. 60 dpp; Immunopositivity is observed in Sertoli cells cytoplasm protrusions from the base to the lumen of the tubules, and in cell-cell junctions. In mature testes, the signal also appears in SPZ. J, K, L. Negative controls for the same time points, obtained by omitting the primary antibody. Scale bars represent 20 μm. PNA: PNA lectin staining. TUB: Tubulin. bg: Background/autofluorescence. nc: Negative controls. For cell-type pointer legend, see table in Fig. 1.

### Expression of PREP and Tubulin in rat and human spermatozoa

The expression of PREP and Tubulin in rat and human SPZ was assessed by Western Blot on protein extracts from epididymal and ejaculated SPZ, respectively (Fig. 7). The data confirmed the presence of the two proteins in male gametes of both species.

**Fig. 7.**
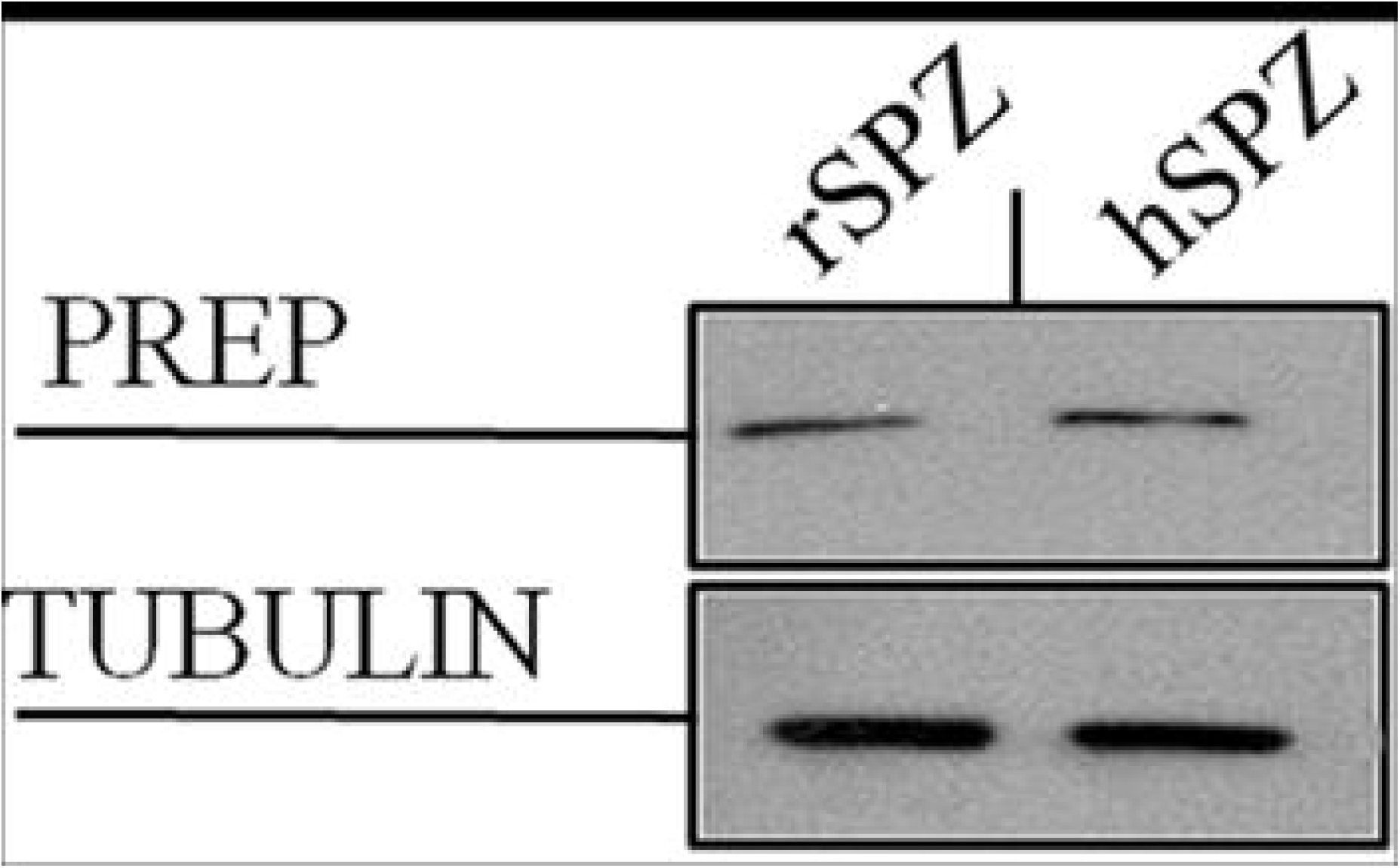
Expression of PREP and Tubulin in rat and human spermatozoa. Western blot analysis on protein extract from rat (lane 1) and human (lane 2) SPZ. PREP (81 KDa, top section) and Tubulin (50 KDa, bottom section) are present in both the samples.

### Co-localization of PREP and Tubulin in rat and human spermatozoa

In order to further expand the profile of PREP localization in male gametes, an immunofluorescence analysis was carried out on rat epididymal SPZ (Fig. 8): there, the protein was mainly detectable inside the flagellum (Fig. 8, C, E), where it clearly co-localize with Tubulin (Fig. 8, D, F). To obtain more detailed data about PREP expression profile in gametes, we extended the analysis on human ejaculated SPZ (Fig. 9). The signal in human gametes confirmed PREP presence inside the flagellum (Fig. 9, C, E), with a weaker signal in the midpiece. Also in this case, PREP and Tubulin co-localize within the flagellum (Fig. 9, D, F), showing a comparable expression pattern described in rat SPZ.

**Fig. 8.**
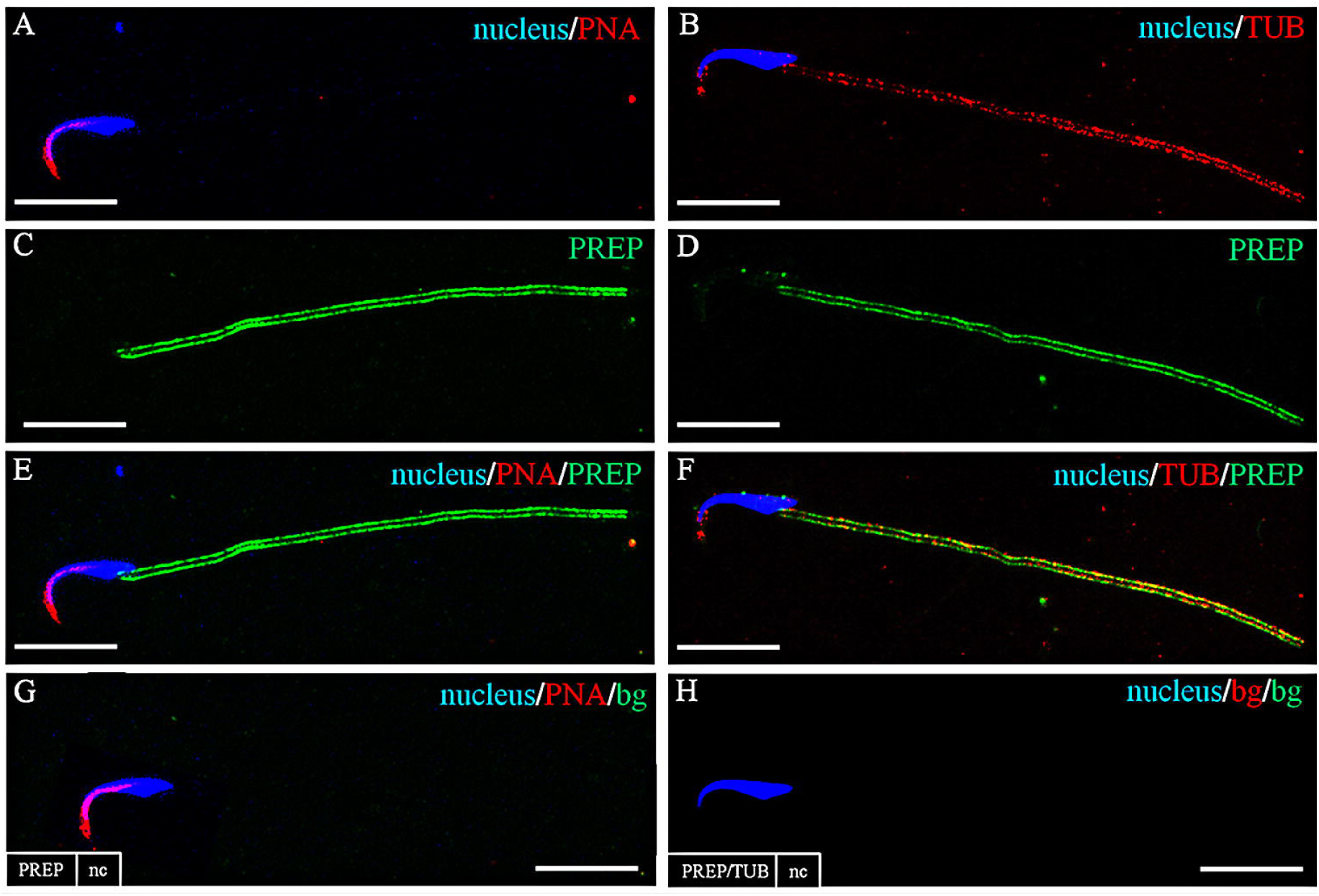
Co-localization of PREP and Tubulin in rat spermatozoa. A: DAPI-fluorescent nuclear staining (blue) and PNA lectin acrosome staining (red). B: Fluorescent signal of Tubulin (red). C-D: Fluorescent signal of PREP (green). E-F: Merged fluorescent channels (blue/red/green) including either PREP or Tubulin, respectively. C-F: PREP is clearly detectable in the flagellum. B: Tubulin marks the region of the tail. F: PREP and Tubulin co-localize inside the flagellum. G-H: Negative controls for PREP or Tubulin, obtained by omitting the primary antibodies. Scale bars represent 10 μm. PNA: PNA lectin staining. bg: Background/autofluorescence.

**Fig. 9.**
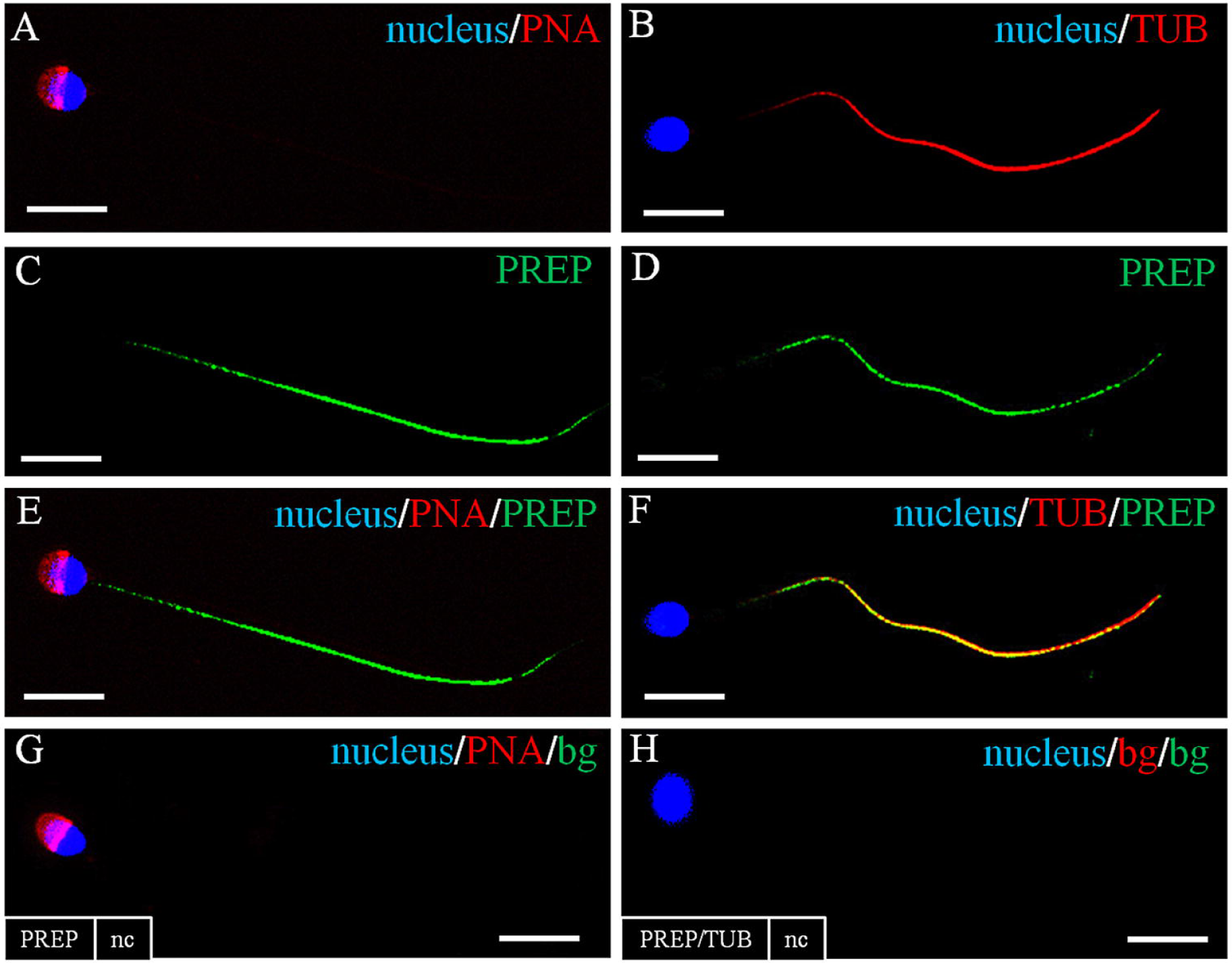
Co-localization of PREP and Tubulin in human spermatozoa. A: DAPI-fluorescent nuclear staining (blue) and PNA lectin acrosome staining (red). B: Fluorescent signal of Tubulin (red). C-D: Fluorescent signal of PREP (green). E-F: Merged fluorescent channels (blue/red/green) including either DAAM1 or Tubulin, respectively. C-E: PREP is clearly detectable in the flagellum, with a weaker signal in the midpiece. B: Tubulin marks the region of the tail. F: PREP and Tubulin co-localize inside the flagellum. G-H: Negative controls for PREP or Tubulin, obtained by omitting the primary antibodies. Scale bars represent 10 μm. PNA: PNA lectin staining. bg: Background/autofluorescence.

## Discussion

In Mammals, the post-natal development of the male gonad is a complex process, during which the seminiferous tubules progressively change their size, structural organization and composition. While the first wave of spermatogenesis takes place germ cells migrate toward the base of the tubule and start their proliferation and differentiation, which will lead to the production of mature spermatozoa (SPZ), while they are nurtured and led by their association with somatic Sertoli cells (SC). Such path is also marked by a significant cytoskeletal remodelling that allows for the formation of complex structures, which allows germ cell separation (GC), protection and maintenance (Pariante et al., 2016).

It is well known that tubulin is one of the key factors involved in these processes, through the regulation of its polymerization and stabilization. Among its many roles, the protein is important in SC for the formation of their wide cytoplasmic protrusions (Lie et al., 2010), which follow differentiating GC toward the lumen. In particular, in SC cytoplasm, the microtubules are orientated in linear arrays parallel to the long axis of the cell (Vogl et al., 1995). Microtubules are evident in the lateral processes surrounding round and elongating spermatids (SPT) (Amlani and Vogl, 1988; Vogl, 1988;) and they show complex changes as germ cells progress through the various stages of seminiferous cycle (Vogl et al., 2008).

It is also known that prolyl endopeptidase (PREP), a serine protease enzyme able to digest small peptides and involved in several physiological and pathological processes, has been associated to microtubules, and in particular with the C-terminus of α-tubulin, suggesting that this endopeptidase may be involved in microtubule-associate processes, independent of its peptidase activity (Schulz et al., 2005). Many studies showed that the protein may have a very important role in the central nervous system (Mentlein, 1988; Wilk, 1983), but it has been also involved in the physiology of other districts, as well as reproductive organs. Indeed, PREP was originally found as an oxytocin-cleaving enzyme in human uterus (Walter et al., 1971), and later implicated in male gametogenesis: it was purified from ascidian sperm (Yokosawa et al., 1983); then the protein was isolated from herring testis (Yoshida et al. 1999). Then, PREP was localized in mouse spermatids and SPZ and it was hypothesized that it may be involved in sperm motility (Kimura et al., 2002). Later analyses on human showed that PREP localizes in the seminiferous tubules and Leydig cells and proposed that it may participate in regulating the levels of seminal TRH analogues, mediating death associated with necrozoospermia (Valdivia et al., 2004; Myöhänen et al., 2012). Finally, in our previous work (Dotolo et al., 2016) we studied the effects of PREP knockdown on testis and sperm in adult mice, showing that the enzyme is indeed needed for a correct reproductive function and that its absence leads to marked alterations of the gonads and, ultimately, gametes. All these reports suggest that PREP might have an active role in male reproductive function. In the present study, in order to improve upon the current knowledge, we investigate the possible association of PREP with the morphogenetic changes which occur during the post-natal development of rat testis, choosing a time frame ranging from 7 to 60 days post-partum, which represents the first wave of spermatogenesis.

The first, encouraging evidence comes from the Western Blot expression data: PREP is, indeed, expressed in the developing and adult testis. The successive localization analysis highlighted that the protein localizes in the cytoplasm of proliferating SC during all the stages of development. It is worthy of note the congruence between PREP profile and tubulin distribution in SC protrusions, which surround the GC. It is well known that GC translocation, and in particular that of SPT, through the seminiferous epithelium occurs via microtubules-based transport of the apical ectoplasmic specialization (ES), a structural connection between SC and differentiating GC (Su et al., 2013). It has been proposed that the microtubules in this process act as a “rail” for the re-localization of cellular contents, as well as of translocation of SPT, which is obtained by the gliding of the entire ES structure together with attached SPT along microtubules within SC (Lie et al., 2010). As said before, PREP has been associated with the C-terminus of α-tubulin, this information, coupled with our result, may suggest a possible involvement of PREP in such cytoskeletal remodelling.

On the other hand, we detected PREP presence inside the proliferating and differentiating GC during the first wave of spermatogenesis. Spermatogonia are immature GC which undergo a series mitosis to give rise to a pool of cells that enter in meiosis: I and then II spermatocites. It has been already reported that PREP inhibition suppressed the growth of human neuroblastoma cell (Matsuda et al., 2013) and that it may be a positive regulator of cell cycle progression in human gastric cancer cell (Suzuki et al., 2014). More recently, PREP was found in various cell types in both the cytoplasm and nuclei in mouse whole-body sections, in co-localization with Ki-67, a proliferation marker protein, suggesting its role in cell proliferation (Myöhänen et al., 2012). Here we can hypothesize that PREP may be involved in the meiotic and post-meiotic phases of GC differentiation. Our supposition is corroborated by the co-localization of this peptidase with tubulin: as well known, microtubules are the main elements of mitotic and meiotic spindles, which help the division of chromosomes/chromatids into the two daughter cells.

During spermiogenesis and in adult testis, PREP co-localize with tubulin in the cytoplasm of haploid round and elongating SPT. This may correlate either with the aforementioned physiological activity of SC, which maintain and hold the spermatids until their release during spermiation, or with spermiohistogenesis itself, during which PREP may be needed for the correct organization and differentiation of SPT. In fact, dynamic microtubules are essential for the assembly of microtubule-based structures that participate in SPT remodelling and physiology, such as the manchette and the sperm flagella. Thus, such wide distribution may hint at a possible function for PREP, due not only of its enzymatic activity, which could led to the degradation and maturation of small active molecules involved in the process, but also with its association with microtubules and its involvement in all microtubules-associated processes which take place during the spermatogenesis. It is interesting to note, that the endopeptidase is clearly detectable in the tail of isolated epididymal and human SPZ, as well as tubulin, which suggests that the PREP may be involved in mature sperm function. Our data, which match with those found in our previous work by Dotolo et al. (2016) in mouse sperm, let us to hypothesize a possible role of the enzyme in mammalian sperm motility: as known, the process is driven by the release and uptake of calcium by intracellular stores (Herrick et al., 2005; Ho and Suarez, 2003), and being PREP a possible regulator of the pathway of inositol 1,4,5 which results in the modulation of cytosolic calcium level (Szeltner and Polgár, 2008) we suggest that PREP, through its involvement in calcium signalling, might be an actor in the regulation of sperm movement and progression. As known, the motility is generated by the internal cytoskeletal structure called axoneme, a highly organized microtubule-based structure constructed from approximately 250 proteins that has been well conserved through evolution (Inaba, 2003). In this case, PREP may have a double function in sperm motility: one regarding the aforementioned modulation of cytosolic calcium level, and the other concerning the regulation of the pivotal role that has the tubulin in the progression of the sperm movement.

Thus, our data strongly support the hypothesis that PREP could be considered as a useful marker in further studies aimed at the observation of the reproductive function and the physiological sperm motility, if enhanced for such purpose, due to its wide distribution in the cytoplasm of SC, GC and in the flagellum of mammalian SPZ.

In conclusion, our work shows the expression and the localization of PREP during rat spermatogenesis and in rat and human SPZ. Although the exact functions of this enzyme remain to be elucidated, it is clear that PREP is involved in spermatogenetic events. The results here described represent a starting point to understand and define the effective role of the endopeptidase in mammalian reproduction, in order to be able to use PREP as a marker of a good quality of the gamete physiology.

The authors declare no conflict of interest

Contract grant sponsor: “Ricerca di Ateneo” Università degli Studi della Campania “Luigi Vanvitelli”.

